# *STX18-AS1* is a Long Noncoding RNA predisposing to Atrial Septal Defect via downregulation of *NKX2-5* in differentiating cardiomyocytes

**DOI:** 10.1101/2020.05.27.118349

**Authors:** Yingjuan Liu, Mun-kit Choy, Sabu Abraham, Gennadiy Tenin, Graeme C. Black, Bernard D. Keavney

## Abstract

Previous genome-wide association studies (GWAS) have identified a region of chromosome 4p16 associated with the risk of Atrial Septal Defect (ASD), which is among the commonest Congenital Heart Disease (CHD) phenotypes. Here, we identify the responsible gene in the region and elucidate disease mechanisms. Linkage disequilibrium in the region, eQTL analyses in human atrial tissues, and spatio-temporal gene expression studies in human embryonic hearts concordantly suggested the long noncoding RNA (lncRNA) *STX18-AS1* as the causative gene in the region. Using CRISPR/Cas9 knockdown in HepG2 cells, *STX18-AS1* was shown to regulate the expression of the key cardiac transcription factor *NKX2-5* via a *trans-*acting effect on promoter histone methylation. Furthermore, *STX18-AS1* knockdown depleted the potential of human embryonic stem cells (H9) to differentiate into cardiomyocytes, without affecting their viability and pluripotency, providing a mechanistic explanation for the clinical association.

---

Congenital heart disease (CHD) is the commonest birth defect, affecting approximately 9.4/1000 live births ^1^. 80% of CHDs behave as complex diseases of multifactorial determination. Genetic investigations of CHD susceptibility have revealed roles for rare single nucleotide polymorphisms and copy number variants, and shown association with common SNPs in GWAS studies ^2–9^.

A region on chromosome 4p16 was among the first GWAS regions identified in CHD, showing association with Atrial Septal Defect (ASD), which is the presence of an abnormal communication between the cardiac atria ^3^. ASD is one of the commonest CHD clinical phenotypes, which frequently requires treatment either in childhood or adulthood, and can lead to serious complications such as right heart failure and cardiac arrhythmia. The 4p16 association was confirmed by subsequent studies and meta-analysis ^10–12^; the causative gene in the region and mechanism of the phenotypic association, however, remain unknown.

In the 4p16 region, linkage disequilibrium around the lead SNPs for ASD is confined to a genomic segment including the long noncoding RNA *STX18-AS1* (Supplementary figure 1). Previously, the most strongly associated SNPs (rs870142, rs6824295 and rs16835979) have been shown to be eQTLs for *STX18-AS1* in adult ventricular myocardial tissue ^3^. These observations suggested further mechanistic investigation of *STX18-AS1* as a candidate gene for ASD. Long noncoding RNAs (lncRNA) are versatile regulators involved in various biological processes; a number of lncRNA genes have been shown to play a role in heart development with *in vitro* or *in vivo* models ^13–15^. Therefore, in this study, we carried out mechanistic investigation of *STX18-AS1* as a candidate gene for ASD. As *STX18-AS1* exhibits poor sequence conservation beyond primates (Supplementary figure 2) all investigations were performed in human cell lines and tissues.

## Results

### Expression studies of *STX18-AS1* in human cardiac tissue

First, we confirmed that the risk SNPs for ASD, located in the second intron of *STX18-AS1*, were tissue-specific eQTLs in 108 human adult atrial tissue samples (Supplementary figure 3 A-E). The SNP genotypes were associated with 59% lower *STX18-AS1* expression per risk allele (p=0.038-0.039) in atrial tissue, with no association with expression in peripheral blood (Supplementary figure 3 F). The finding in human atrial tissue therefore confirms the previous report in human ventricular tissue ^3^.

We then quantified the expression of *STX18-AS1* in human embryonic tissues at different developmental stages (CS, Carnegie stages). By comparison to adult tissues, *STX18-AS1* was enriched in foetal tissues, including heart, brain, kidney, liver and lung (Figure 1 A), consistent with a role in development. During heart development, the highest expression of *STX18-AS1* in embryonic hearts was found at CS14-18 (Figure 1 B), which is the critical time period for atrial septation ^16^. After CS18, the cardiac *STX18-AS1* expression was gradually reduced as development proceeded (Figure 1 B). To show the location of *STX18-AS1* expression in embryonic hearts, *in situ* hybridization with *STX18-AS1* probe was performed on whole embryonic hearts of CS17-19. At intracellular level, *STX18-AS1* was confirmed to be a nucleus retained lncRNA (Supplementary figure 4). In whole hearts, *STX18-AS1* expression was detected in the atrial septum from CS17 to CS19 (Figure 2 A-C; N=3 hearts). Sections between CS17 to CS19 confirmed *STX18-AS1* positive cells located within the atrial septal myocardium (Figure 2 D-F; N=3 hearts). In addition to the atrial septum, the atrioventricular canal (AVC), part of the outflow tract (OFT) and ventricles were also stained (Figure 2 A). However, comparing with the other parts of the embryonic heart at CS15, the relative quantity of *STX18-AS1* transcripts in the atrial septum, assessed by qPCR, was the highest (Supplementary figure 5). Thus, *STX18-AS1* expression shows strong spatio-temporal correlation with atrial septation during human heart development.

**Figure 1:**
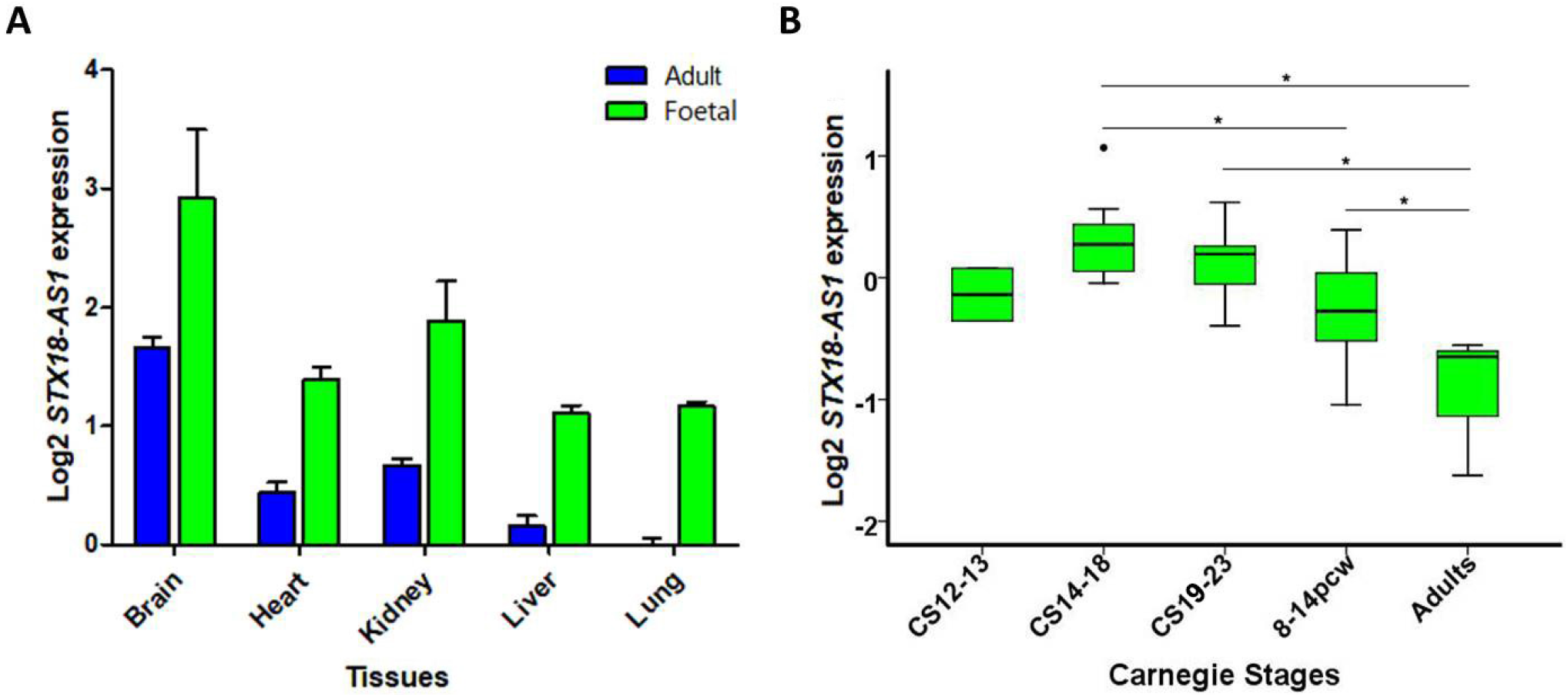
Expression of *STX18-AS1* in human tissues. **A**, *STX18-AS1* expression in different tissues from adults and foetus. **B**, expression of *STX18-AS1* in adult and developing hearts of different Carnegie stages (CS). Except for stage CS12-13 (2 samples), ≥ 3 bio-replicates were done for each group. Data are shown as Mean±SD in A, and distributions in B. Difference between groups was tested by ANOVA. * indicates *P*<0.05.

**Figure 2:**
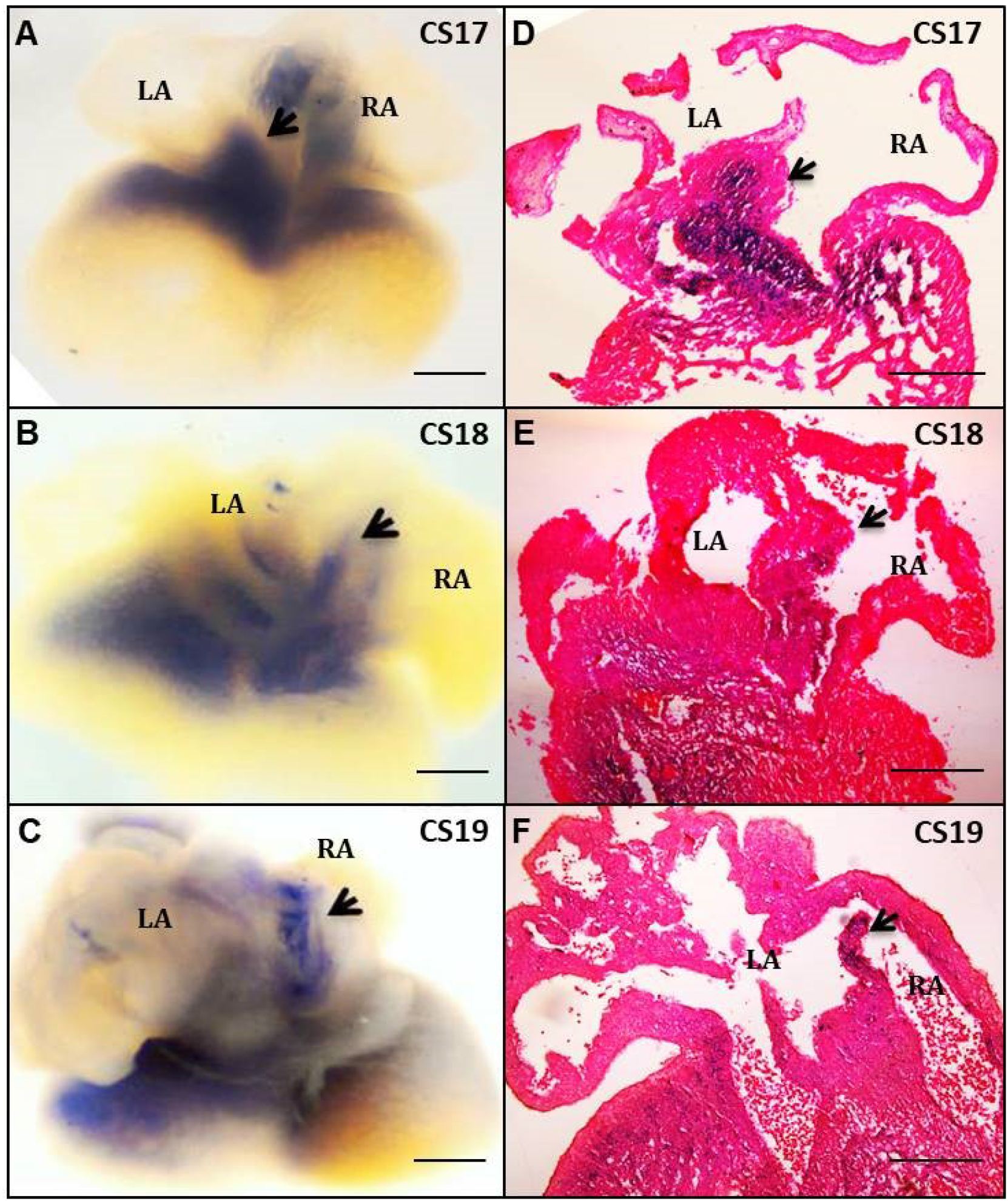
In situ hybridisation on human embryonic hearts with *STX18-AS1* probe. **A-C**, whole embryonic hearts of CS17-19 hybridised with *STX18-AS1* antisense probes. **D-F**, embryonic heart sections stained with Eosin Red. Blue colour indicates the expression of *STX18-AS1* in hearts. Black arrows refer to the location of Atrial Septum. RA, Right Atrium; LA, Left Atrium. Scale bars represent 200μm.

### *STX18-AS1* regulates the expression of the key cardiac transcription factor *NKX2-5*

Since the risk genotypes at GWAS SNPs for ASD were associated with lower *STX18-AS1* expression, we used CRISPR/Cas9 to analyse the downstream signalling consequences of *STX18-AS1* knockdown. It is reported that lncRNAs are more resistant to point mutations than protein-coding genes, and that large deletions may be necessary to knock down the transcription of lncRNAs ^17^. We therefore adopted a strategy to delete the first two exons of *STX18-AS1.* Two pairs of gRNAs were designed (STX18-AS1 CRISPR and STX18-AS1 CRISPR 2^nd^; Supplementary figure 6 A); results of the first design (STX18-AS1 CRISPR) are reported in the main text and results of the second design (STX18-AS1 CRISPR 2^nd^) in supplementary data. Observations for both CRISPR pairs were concordant. Control cells were transduced with empty CRISPR vectors without gRNA (Control CRISPR).

We applied CRISPR in HepG2 cells as we established these cells have readily detectable expression of *STX18-AS1*, and the key cardiac transcription factors *NKX2-5*, *GATA4*, and *MSX1* (Supplementary figure 7). With both CRISPR designs, the genomic deletions of the first two exons of *STX18-AS1* was achieved, producing a short product of ~600bp in compare to the original sequence of 3.3kb and 4.1kb (Supplementary figure 6 B-E). The sequence of the short products was well aligned to the genome except for the junctions at gRNA targeting sites (Supplementary figure 8 A-B). The STX18-AS1 CRISPR decreased the transcription of *STX18-AS1* in HepG2 by ~70% (Figure 3 A). Transcription of *NKX2-5* was reduced by ~80% by knockdown of *STX18-AS1* (Figure 3 B-D); the protein production of *NKX2-5* in HepG2 cells was also inhibited by *STX18-AS1* knockdown with no effect on the reference gene β-actin (Figure 3 E-I). The same effect on *NKX2-5* expression was achieved with STX18-AS1 CRISPR 2^nd^ (Supplementary figure 9 A-G). Therefore, the CRISPR knockdown experiments of *STX18-AS1* suggested that *NKX2-5* was a downstream target of *STX18-AS1*.

**Figure 3:**
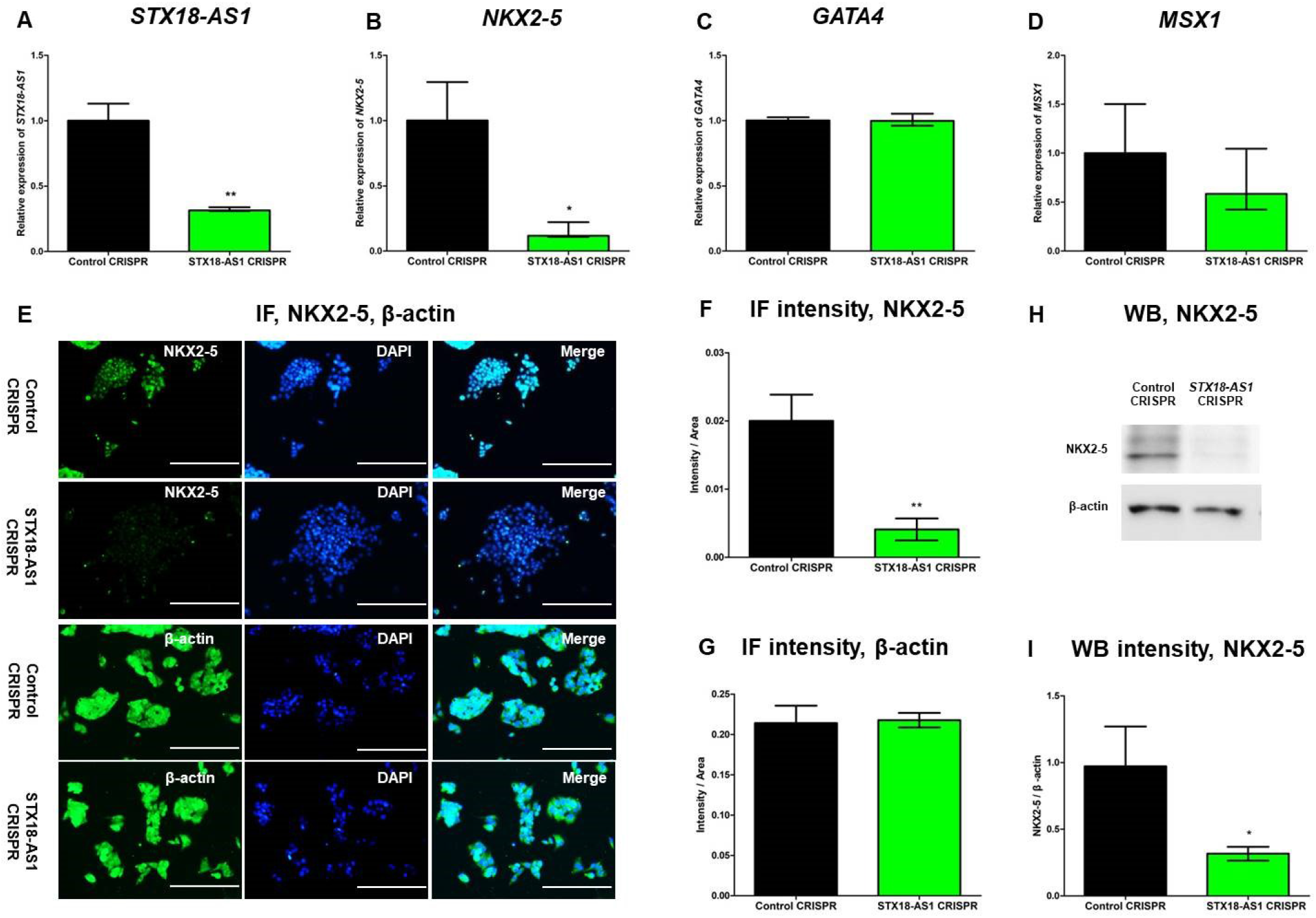
CRISPR knockdown of *STX18-AS1* inhibits the expression of *NKX2-5* in HepG2 cells. With CRISPR knockdown, transcription of *STX18-AS1* was reduced **(A)**, and the level of *NKX2-5* transcripts decreased **(B)** without changes in *GATA4* **(C)** and *MSX1* **(D)** mRNA. The protein level of NKX2-5 was reduced with reference to β-actin, detected with both immunofluorescence **(E)** and immunoblotting **(H).** The quantitative intensity analyses confirmed the statistical difference between Control CRISPR and STX18-AS1 CRISPR for both immunofluorescence **(F-G)** and immunoblotting **(I).** Data are shown as Mean±S.E. *, *P*<0.05; **, *P*<0.01, comparing with Control CRISPR. Scale bars represent 200μm.

### *STX18-AS1* reduces *NKX2-5* expression via effects on chromatin remodeling

To further investigate the potential mechanism of *STX18-AS1* in ASD, epigenetic changes surrounding *NKX2-5* consequent on *STX18-AS1* knockdown, and the interaction of *STX18-AS1* with chromatin remodeling complexes were evaluated in HepG2 cells. Consequent upon STX18-AS1 CRISPR knockdown, the histone marker indicating an active promoter (H3K4me3) around *NKX2-5* was decreased by ~70%, commensurate with the reduction of *NKX2-5* expression, without changes in its nearby repressive marker H3K27me3 (Figure 4 A). Similar results were also achieved with the STX18-AS1 CRISPR 2^nd^ (Supplementary figure 9 H). We then used Comprehensive Identification of RNA-binding Proteins (ChIRP) ^18^ to investigate the interaction between *STX18-AS1* and WDR5, the scaffold protein of the SET1/MLL complex. The function of the SET1/MLL complex is necessary for the trimethylation of H3K4^19^. The interaction between U1 snRNA and U1A protein ^20^ served as the positive control (Figure 4 B). *STX18-AS1* transcripts were successfully retrieved by *STX18-AS1* probes (Figure 4 C). Along with the retrieved *STX18-AS1* transcripts, WDR5 was detected in the protein pulldown elution, with no signal in the negative control or background control (β-actin) (Figure 4 D). To demonstrate the specificity of the interaction, we also quantified interaction between *STX18-AS1* and SUZ12, a core subunit of PRC2 complex catalyzing the methylation of H3K27 ^21^. No signal of SUZ12 interaction was detected in the pulldown protein sample (Supplementary figure 10). These results indicate *STX18-AS1* interacts with WDR5 protein with a degree of specificity, linking levels of *STX18-AS1* to the mechanism of H3K4 trimethylation.

**Figure 4:**
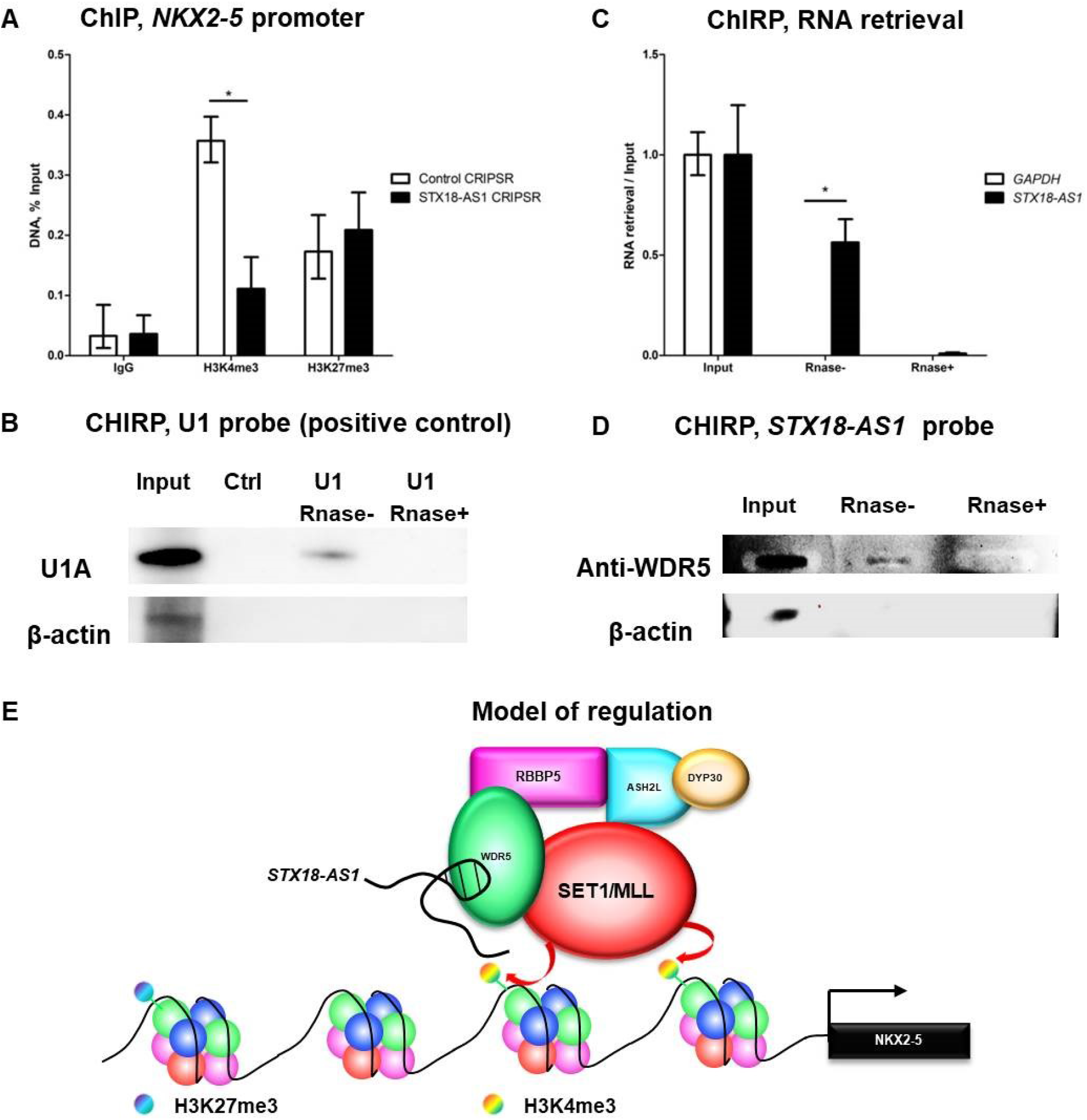
*STX18-AS1* regulates *NKX2-5* in *trans* through histone methylation. **A**, Using ChIP, H3K4me3 around the promoter of *NKX2-5* was reduced after CRISPR knockdown of *STX18-AS1*, without changes in H3K27me3. IgG was used as a background control. **B**, ChIRP was tested with U1 probe (interacting with U1A protein) as a positive control. Ctrl, random probe negative control; Rnase+, RNase treated negative control; β -actin was a negative control for protein precipitation. **C**, The RNA of *STX18-AS1* was successfully retrieved with *STX18-AS1* ChIRP probes. GAPDH represents the genetic background control. **D**, WDR5 protein (detected with slot blotting), the scaffold protein of MLL complex (which modifies H3K4me3), was pulled down by *STX18-AS1* probes, but the signal was absent on RNase treatment (Rnase+). **E**, the model of epigenetic regulation around *NKX2-5* by *STX18-AS1*. Data are shown as Mean±S.E. *, *P*<0.05.

### *STX18-AS1* knockdown reduces *in vitro* cardiomyocyte differentiation

It is known that mutations in *NKX2-5* are causative of septal defects in human families ^22^. We next investigated whether *STX18-AS1* knockdown affected the differentiation from human embryonic stem (ES) cells to cardiomyocytes, a process in which *NKX2-5* is known to play a critical role. We used the H9 human ES cell line (H9) in view of established protocols directing differentiation of these cells into beating cardiomyocytes ^23,24^. The genomic deletion at *STX18-AS1* was achieved in H9 cells with the same STX18-AS1 CRISPR strategy, generating a ~50% decrease in *STX18-AS1* transcription (Supplementary figure 11 A-B). No obvious change in cell morphology (Supplementary figure 11 C), proliferation (Supplementary figure 11 D), apoptosis rate (Supplementary figure 11 E-F) and pluripotency (Supplementary figure 12) in the naïve state of H9 cells was observed during maintenance culture after *STX18-AS1* knockdown.

Following the *STX18-AS1* knockdown in H9 cells, the differentiation potential into cardiac lineage was examined following a protocol developed from Lian et al.’s method ^23^. In control cells, *STX18-AS1* transcription increased from Day 6 onward, while cells targeted with the STX18-AS1 CRISPR did not (Figure 5 A). Correspondingly, no substantial differences in *NKX2-5* transcription (Figure 5 B) were evident before Day 6. After Day 6, reduced beating cell colonies (online supplementary video) and decreased *NKX2-5* expression (Figure 5 B) in STX18-AS1 CRISPR H9 were observed. At Day 8, beating cells were identified in 90% wells (18/20) of Control CRISPR H9 cells, while the rate for STX18-AS1 CRISPR H9 cells was only ~20% (4/23; *P*<0.001). Fewer beating cells differentiated from STX18-AS1 CRISPR H9 cells, compared to Control CRISPR H9 cells, in which the majority of cells were beating robustly (online supplementary video: DOI: http://dx.doi.org/10.17632/vdf4nyfd9y.2#file-b13de0de-52eb-487e-978b-c85d79fe83ac & DOI: http://dx.doi.org/10.17632/vdf4nyfd9y.2#file-1b02d4db-8b98-4f02-803a-1dc15145b8b1); there was a corresponding reduction in the expression of Cardiac Troponin T (C-troponin T) and the proportion of C-troponin T positively stained cells (Figure 5 C, Supplementary figure 13). These observations suggest a potential mechanism whereby atrial septal defects could result from genetically determined lower levels of *STX18-AS1* during human cardiac development.

**Figure 5:**
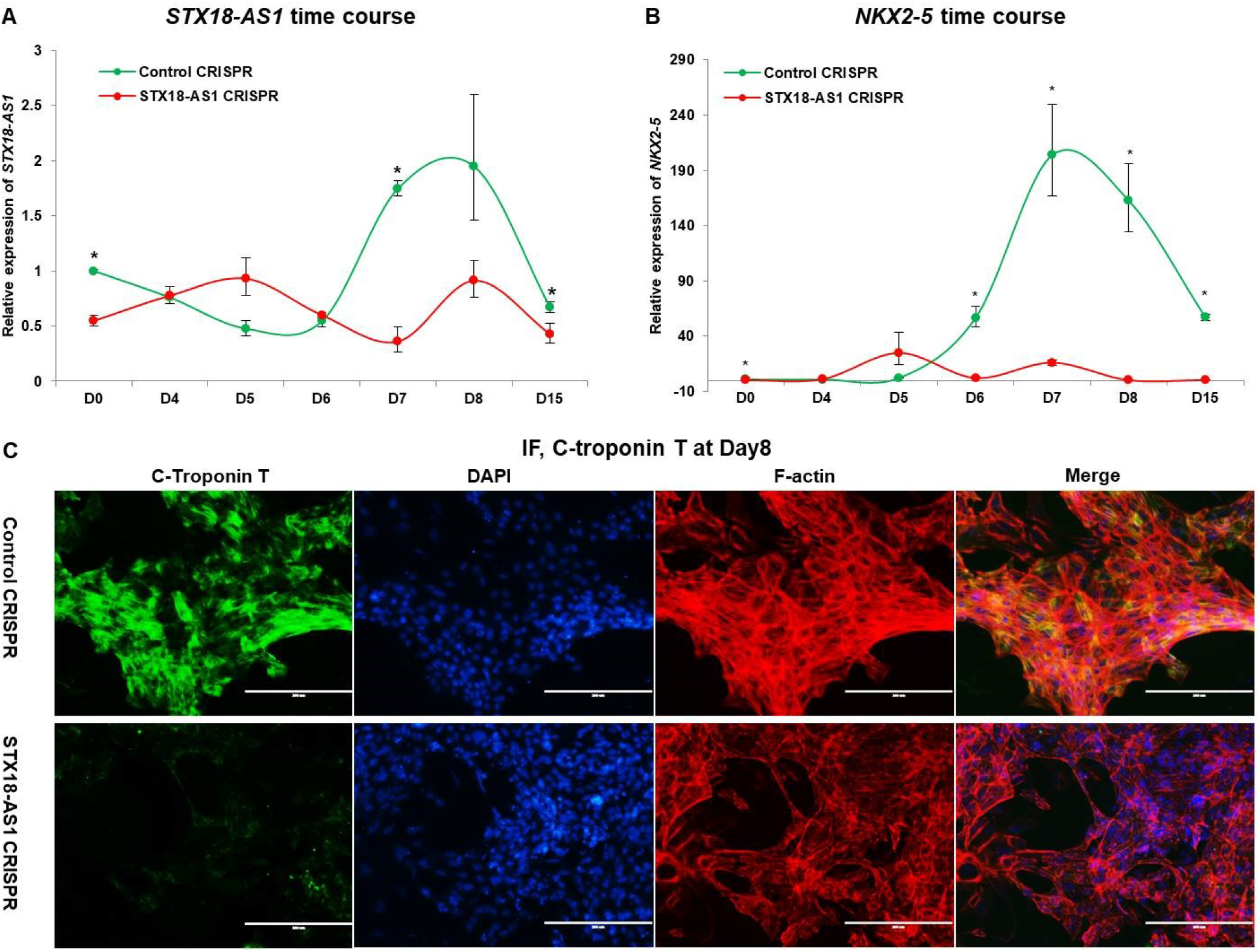
Knockdown of *STX18-AS1* depletes the potential of human embryonic stem cell (H9) for cardiomyocyte differentiation. During the *in vitro* cardiomyocyte differentiation protocol, the expression of *STX18-AS1* **(A)** and *NKX2-5* **(B)** changed dynamically with an peak after D6 for Control CRISPR H9 cells but not *STX18-AS1* CRISPR H9 cells. Beating cardiomyocytes were successfully differentiated in Control CRISPR H9 cells with positive staining of cardiac troponin T (C-troponin T) reference to DAPI and F-actin indicated by Phalloidin **(C).** Beating cells differentiated from STX18-AS1 CRISPR H9 cells were relatively less with weak C-troponin T staining **(C).** Data are shown as Mean±S.E. *, *P*<0.05, comparing between Control CRISPR and STX18-AS1 CRISPR. “D0-D15” refers to the days of cardiomyocyte differentiation.

## Discussion

We present evidence that the lncRNA *STX18-AS1* is the causal gene for ASD susceptibility at chromosome 4p16. Data from meta-analyses of GWAS studies is newly supported by eQTL data in adult human atrium, strong spatio-temporal correlation of *STX18-AS1* expression with atrial septal development in fetal human hearts, and by the demonstration of *trans-*acting regulatory effects on *NKX2-5*, a key septal development gene. Knockdown of *STX18-AS1* inhibits or delays *in vitro* cardiomyocyte differentiation, a plausible mechanism for the phenotypic association between SNPs in the gene - whose risk alleles reduce expression levels - and ASD. *STX18-AS1* is to the best of our knowledge the first lncRNA with a function in cardiac development identified from GWAS.

Through knockdown experiments, we showed that *STX18-AS1* regulates the downstream expression of *NKX2-5* (and possibly other cardiac developmental genes) through chromatin remodelling mediated at least in part via its interaction with WDR5 (Figure 4 E). Such epigenetic mechanisms are recognised for other lncRNAs ^25^.For example, similar to *STX18-AS1*, the previously reported lncRNA *Bvht*, a mouse lncRNA impacting on the initiation of the cardiac lineage, can switch on the cardiac core transcriptional factor network through epigenetic modifications ^26^. *STX18-AS1* is not conserved beyond primates, and the plausibility of a non-conserved gene having a key role in human cardiac development may reasonably be questioned; however, we note that the conservation of *Bvht* across species is if anything even less than *STX18-AS1*, having no homolog in rat or human ^27^. It is known that sequence conservation of lncRNAs is not necessarily a predictable indicator of biological function, with changes impacting secondary structure having particular importance in determining RNA-protein interactions.^28^ Further investigations will be needed to establish whether lncRNAs with similar secondary structures to *STX18-AS1* but lacking sequence homology play cognate roles in heart development in other species. In the current work, the lack of conservation of *STX18-AS1* restricted our investigation to human cells and tissues.

Our expression data showed that *STX18-AS1* expression is not confined to the atrial septum during development; although the GWAS evidence to date is strongest for ASD, smaller effects of the ASD-associated SNPs, or other genetic variants in *STX18-AS1*, on other CHD phenotypes (for example, ventricular septal defect) are not ruled out, and may emerge as larger numbers of CHD patients are studied in GWAS. Both for Mendelian syndromic alleles (*e.g., JAG1* mutations causing Tetralogy of Fallot in the context of Alagille syndrome), and susceptibility variants (*e.g.*, SNPs in GATA4 and bicuspid aortic valve), a degree of phenotypic specificity is a typical observation in CHD, to which our findings correspond. Having identified an effect of *STX18-AS1* on the expression of the key septal development gene *NKX2-5*, we pursued this mechanism in our functional studies; effects of *STX18-AS1* on other important transcriptional circuits during development may remain to be discovered.

To date, over 3000 SNP associations have been identified with cardiovascular diseases in the NHGRI GWAS catalog, although causative mechanisms have been identified at only a few loci ^29^. Among all disease-associated SNPs, 45% map to lncRNAs, suggesting an important role for these genes in complex disease susceptibility ^30^. Relatively few annotated lncRNAs containing GWAS associated SNPs for cardiovascular diseases have been investigated ^31^. Recently, the human homolog of the mouse lncRNA *Upperhand* was found to be a control on expression of the cardiac transcription factor *Hand2* and essential for mouse heart development ^32^. Our study of *STX18-AS1* also illustrates the potential for lncRNA studies in human CHD. Therefore, in future work, investigation of common human genetic variation in other lncRNA genes would be of considerable interest.

Maximising the differentiation and maturation of stem-cell derived cardiomyocytes is a key challenge for myocardial regenerative medicine strategies. Investigations of the cardiomyocyte differentiation processes are therefore of potential clinical importance. Since the risk alleles for ASD caused a reduction in *STX18-AS1* expression, we focused on knockdown experiments in our mechanistic investigations. Knockdown of *STX18-AS1* markedly inhibited cardiomyocyte differentiation from hES cells. It would be of interest to determine whether overexpression of *STX18-AS1* at an appropriate time in differentiation could enhance the differentiation of ES-cell derived cardiomyocytes, or stimulate their development towards more mature and clinically useful phenotypes.

## Methods

### Human tissues

For the expression study, the pooled cDNA samples of human foetal tissues (Fetal MTC Panel, 636747, Lot#1308272A) and adult tissues (MTC panel I, 636742, Lot#1303120A) were sourced from Clontech. The cDNA samples from whole human embryonic and foetal hearts of different developing stages were obtained from the MRC-Wellcome Human Developmental Biology Resource (HDBR). For eQTL analyses, RNA from 108 right atrial appendage (RAA) samples and DNA from the corresponding human blood samples were extracted as previously described^33^. For *in situ* hybridisation, human embryonic hearts fixed in 4% PFA were also obtained from the MRC-Wellcome HDBR.

### CRISPR/Cas9 and cell culture

Two pairs of gRNAs (Table S1 in “Supplementary file”) intermediated by human U6 (hU6) promoter were cloned after hU6 promoter of the plentiCRISPR_V2 plasmid (Addgene, #52961), according to Vidigal *et al.*’s protocol ^34^. The plasmids were packaged into lentivirus by co-transfecting HEK293T cells with pPAX2 (Addgene, #35002) and pMD2.G (Addgene, #12259) in a proportion of 4:3:2 using Lipofectamine 2000 (ThermoFisher). HEK293T cells were maintained in DMEM complete medium (DMEM [Gibco] supplemented with 10% of FBS and 100 UI of Penicillin/Streptomycin). After transfection, the medium of the first 12 hours was discarded and changed into fresh DMEM complete medium. The lentivirus was collected after an additional 48 hours for immediate use in transduction or aliquoted and stored in −80°C for future use.

HepG2 cells were maintained with DMEM complete medium. Lentivirus transduction on HepG2 cells was conducted at the confluency of ~60%, by culturing in Lentivirus soup (with 10μg/ml polybrene [Millipore]) for 24 hours before puromycine selection (in 1μg/ml puromycine [ThermoFisher]). Cells were deubiquitylated with MG132 (10μM) for 24 hours before protein detection with immunofluorescence and immunoblotting.

H9 (human embryonic stem cell line [WiCell]) was maintained with mTesR1 medium (STEMCELL Technologies) supplemented with 10μM ROCK inhibitor (Millipore) in Matrigel-coated plates. Triple lentivirus transductions (refreshed with new lentivirus soup once a day for three days) on H9 cells (Passage 35) were conducted before puromycine selection. The first transduction started at the confluency of 40-60%. For each transduction, cells were permeablised first with 1ml/well (6-well plate) mTESR1 medium with 8μg/ml polybrene (Millipore) for 15 min (37°C). Afterwards, 2ml lentivirus soup with 8μg/ml polybrene was added (24 hours). After the 3^rd^ transduction, maintenance in mTesR1 medium for 1-2 days was required before puromycine selection (0.8μg/ml puromycine [ThermoFisher] for 24 hours). 24-hour puromycine selection was made for every passage.

### Directed cardiomyocyte differentiation

Cardiomyocyte differentiation was performed as previously described ^23^ and summarised as follows. The differentiation was started (Day 0) on H9 monolayers at the confluency of 90-100%. During Day 0-2, H9 cells were cultured with 6μM Chir99021 (GST inhibitor [Millipore]) in B27-medium (RPMI 1640 medium [Invitrogen] supplemented with 1× B27 minus insulin [50×, Gibco]). At Day 2-4, cells were treated with 2μM C59 (WNT antagonist [Abcam]) in B27− medium. Day 4-8, cells were maintained in B27− medium. All mediums were refreshed every day. Since Day 8, cells were maintained with B27+ medium (1×B27 [50×, Gibco] supplemented RPMI 1640 medium [Invitrogen]) till Day15. Beating cells were typically seen since Day 6-8.

### Polymerase chain reaction (PCR)

Genomic DNA was extracted with PureLink Genomic DNA Mini Kit (Invitrogen). PCR primers (Table S2 in “Supplementary file”) for genotyping after CRISPR application were designed using Primer 3 (primer3.ut.ee/). PCR was conducted using OneTaq 2x Master Mix (New England Biolabs) with forward and reverse primers following the program of 95°C×10min, 40× (95°C×20s, 58°C×30s, 68°C×3min), 68°C×10min. The sequences of products were checked by Sanger sequencing (service provided by Genomic Technologies Core Facility in the University of Manchester).

### Quantitative polymerase chain reaction (PCR)

RNA from HepG2 cells were extracted using RNAqueous-Micro Kit (Ambion), while RNA from H9 cells were extracted with Trizol (Invitrogen) following the manufacture’s protocol. The first cDNA strand was synthesised from RNA using M-MLV Reverse Transcriptase (Promega) with both random hexamer and oligo(dT) primers (Promega) after the DNase (Promega) treatment. Gene expression was quantified by TaqMan assays (ThermoFisher, Table S3 in “Supplementary file”) using ViiA7 qPCR system (Life technology). For each qPCR reaction, ~40ng cDNA templates were used with TaqMan™ Gene Expression Master Mix (2×) and TaqMan assays (20×). Three replicates were done for each reaction with the program of 95°C×10min, 40× (95°C×30s, 60°C×1min). *IPO8* was included as the internal control for expressional quantification. For genotyping, Taqman SNP genotyping assays (ThermoFisher, Table S3 in “Supplementary file”) were used for the detection of genomic DNA samples.

### *In situ* hybridisation

Whole-mount In situ hybridisation (ISH) was conducted as previously described ^35^. The DNA templates used for DIG-labeled *STX18-AS1* RNA probe *in vitro* syntheses (Supplementary file) with T3 RNA polymerase (Promega) were amplified from the coding sequence of *STX18-AS1* from human foetal hearts cDNA (obtained from HDBR). Histological sections were produced using the method described in “Supplementary file”.

### eQTL analyses

*STX18-AS1* expression analyses and SNP genotyping in human samples were done with qPCR as described above. The data were analysed with SPSS with a linear regression model on the expression of *STX18-AS1* and the genotypes of SNPs.

### Immunofluorescence (IF)

Cells were fixed with 4% PFA for 15min at room temperature (RT). After three PBT (PBS with 0.5% Triton-100) washes, cells were blocked with blocking buffer (PBT with 10% sheep serum) for 30min at RT. Afterwards, cells were incubated with primary antibody (diluted in PBT with 1% sheep serum) overnight at 4°C. In the next day, cells were incubated with fluorophore-conjugated secondary antibodies (1 hour at RT in the dark) after PBT washes. Following additional washes, cells were incubated with DAPI (1:1000 [Invitrogen] in PBS) and AF680-Phalloidin (for F-actin, 1:1000 [Invitrogen]) for 20min at RT. After three PBT washes, cells were photographed using a fluorescence microscope (EVOS FL). The intensity of fluorescent signal was analysed with ImageJ ^36^. Primary antibodies used for IF were NKX2-5 (mouse, R&D, 1:100), c-Troponin (rabbit, Abcam, 1:200), β-actin (rabbit, Invitrogen, 1:100). Secondary antibodies used for IF were goat-anti-mouse H&L FITC (Cohesion Biosciences) or goat-anti-rabbit H&L FITC (Cohesion Biosciences), with a dilution of 1:200.

### Immunoblotting

Cells were homogenized by a syringe in cold RIPA lysis buffer (25mM Tris-Cl pH7.5, 150mM NaCl, 1% NP-40, 0.5% Na-deoxycholate, 0.1% SDS, 1× protease inhibitor, and 1 mM PMSF). Collected protein samples were boiled with 1× Laemmli buffer before SDS-PAGE procedure with Mini-PROTEAN TGX Precast Gel (Bio-Rad) in Tris/Glycine Buffer (Bio-Rad). After gel running, proteins in the gel were transferred to PVDF membrane (Bio-Rad) with Trans-Blot Turbo Transfer System (Bio-Rad). The blots were incubated with primary antibodies after blocking with 5% milk for 2 hours at RT. Antibodies were NKX2-5 (mouse, R&D, 1:500) and β -actin (rabbit, Invitrogen, 1:1000). Subsequently, the blots were incubated in HRP goat-anti-rabbit or goat-anti-mouse secondary antibodies (Invitrogen, 1:1000) for 2 hours at RT. Finally, the blots were imaged with ChemiDoc™ Imaging Systems (Bio-Rad) using Piece ECL western blotting substrate (Thermo Scientific), or SuperSignal West Pico Chemiluminescent Substrate (Thermo Scientific).

### Chromatin immunoprecipitation-quantitative PCR (ChIP-qPCR)

Cells were crosslinked with 1% formaldehyde for 10min before chromatin fragmentation with EZ-enzyme kit 17-375 (Millipore) following the manufacture’s protocol. For each reaction, 10^^6^ cells were used to prepare a chromatin sample with DNA fragmented into 100-600bp. The ChIP procedure was conducted following Dahl & Collas’s protocol ^37^. In summary, antibodies of IgG (negative control, Abcam, ab2410), H3K4me3 (Abcam, ab8580) and H3K27me3 (Millipore, 07-449) were conjugated to magnetic beads (Dynabeads protein A, ThermoFisher) for chromatin precipitation. DNA was purified from chromatin elutes with Phenol-chloroform-isoamyl alcohol (Invitrogen) and quantified with qPCR (TaqMan custom assay AJOIXG1, targeting at hg19 chr5:172661507-172662070).

### ChIRP-blotting

Following Chu *et al.*’s protocols ^18,38^, 15 biotin-labelled probes targeting *STX18-AS1* RNA were designed (designs in Table S4 in “Supplementary”) and manufactured by 2BScientific. Biotin-labelled U1 probe (positive control) and random probe (negative control) were adapted from Chu *et al.*’s paper ^18^ (sequences in “Supplementary file”) and manufactured by Eurogentec. The method is summarised as follows. Fragmented chromatin lysates (100-500bp) from 100 million crosslinked HepG2 cells were prepared with Bioruptor (Diagnostic). The chromatin lysates (1ml per 100mg cells) were precleared with C-1 magnetic beads (Dynabeads Myone streptavidin C1 [Invitrogen]) before incubation with probes. 100pmol probes were required for incubation with 1ml chromatin lysate at 37°C with rotation overnight. The lysate pretreated with RNase (RNase+ [10μg/ml, Roche]) was used as negative control (RNase+ control). After precipitation with C-1 magnetic beads (100ul beads / 1ml lysates, 37°C×30min), 10% of beads proceeded to RNA extraction with Trizol method to test the efficiency of the RNA pulled down, while the other 90% beads were used for protein purification with TCA precipitation (method in “Supplementary file”). cDNA was reverse transcribed from RNA and quantified with qPCR for *STX18-AS1* (*GAPDH* was used as the genetic background control). Proteins were detected with bis-tris western blotting for U1A (rabbit, Abcam, 1:200), and slot blotting for WDR5 (rabbit, BethyL, 1:1000). For bis-tris western blotting, NuPAGE™ 10% Bis-Tris Protein Gels (ThermoFisher) were used with NuPAGE MOPS SDS Running Buffer (ThermoFisher). For slot blotting, proteins were added directly onto the nitrocellulose membrane (Bio-Rad) with Slot Blotter (CSL-S48 [Cleaver Scientific]).

### Cellular phenotyping

Cell Growth Detection Kit (MTT) (Sigma) was used for detecting cell proliferation, while Alkaline Phosphatase Assay Kit (ALP) (Sigma) was used for evaluating the pluripotency of H9 cells following the protocols of the manufacture. FACS analyses of H9 cell pluripotency were documented in “Supplementary file”. The apoptosis of H9 cells was quantified by double staining with Hochest33342 (for live cells, [ThermoFisher]) and DRAQ7 (for dead cells, [Biostatus]). Pictures were taken with a fluorescence microscope (EVOS FL), and cell numbers were calculated with ImageJ ^36^.

## Acknowledgements

This work was supported by The University of Manchester-Peking University Health Science Centre Alliance and China Scholarships Council. BK holds a British Heart Foundation Personal Chair. We are thankful to Dr Ruairidh Martin for collection of the samples for eQTL analyses and the data of peripheral blood. We thank HDBR group for supplying human hearts samples for expressional analyses.

## Author contributions

YL and BK developed the ideas for this project. YL conducted all the experiments and produced the results. BK and GB supervised on the progress of the project. MC supported on all experiment designs and production. SA contributed to setting up the methods of lentivirus transduction and immunoblotting. GT helped with the *in situ* hybridisation method. YL and BK wrote the manuscripts with contributions from all authors, who have discussed and agreed on the manuscript.

