## Supplementary figures for "*STX18-AS1* is a Long Noncoding RNA predisposing to Atrial Septal Defect via downregulation of *NKX2-5* in differentiating cardiomyocytes"

**Supplementary Figure legends:**

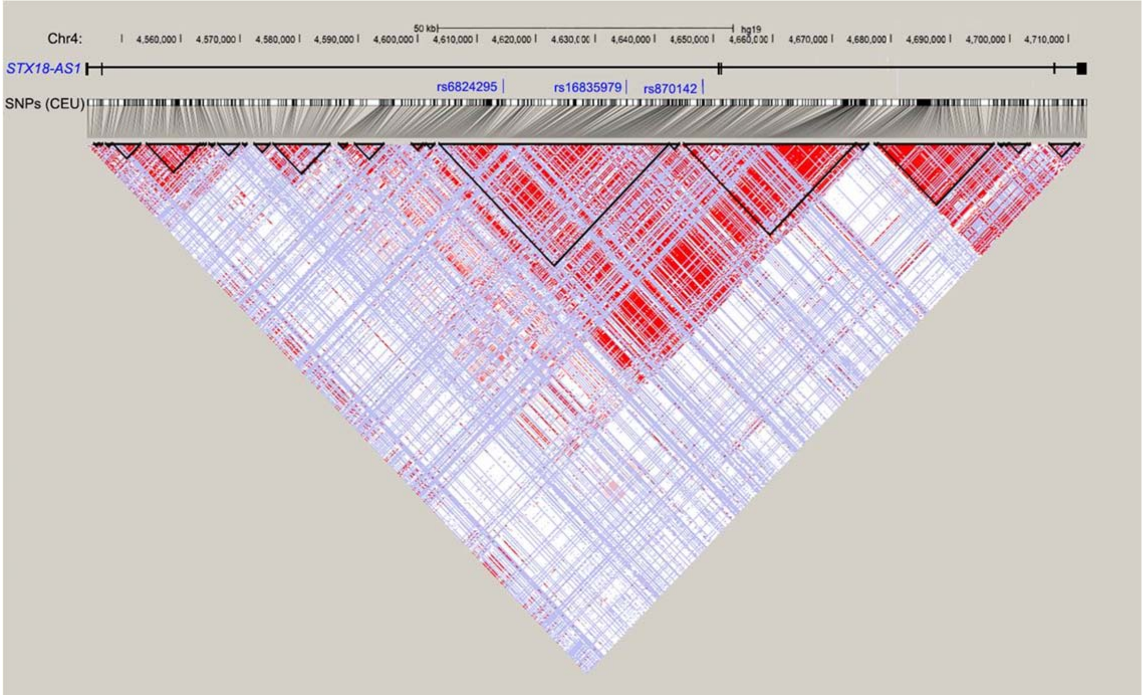


**Supplementary figure 1: Linkage disequilibrium (LD) relationships of SNPs within *STX18-AS1*.**

Data for SNPs in the CEU population was extracted from the 1000 Genomes Project. The LD map was generated with HaploView [1]. The LD of SNP pairs is represented as $r^2$ in white to red (0-1). The locations of the three risk SNPs identified in GWAS are labelled in blue. The black triangles represent haplotype blocks (defining with confidence intervals according to Gabriel's method [2])

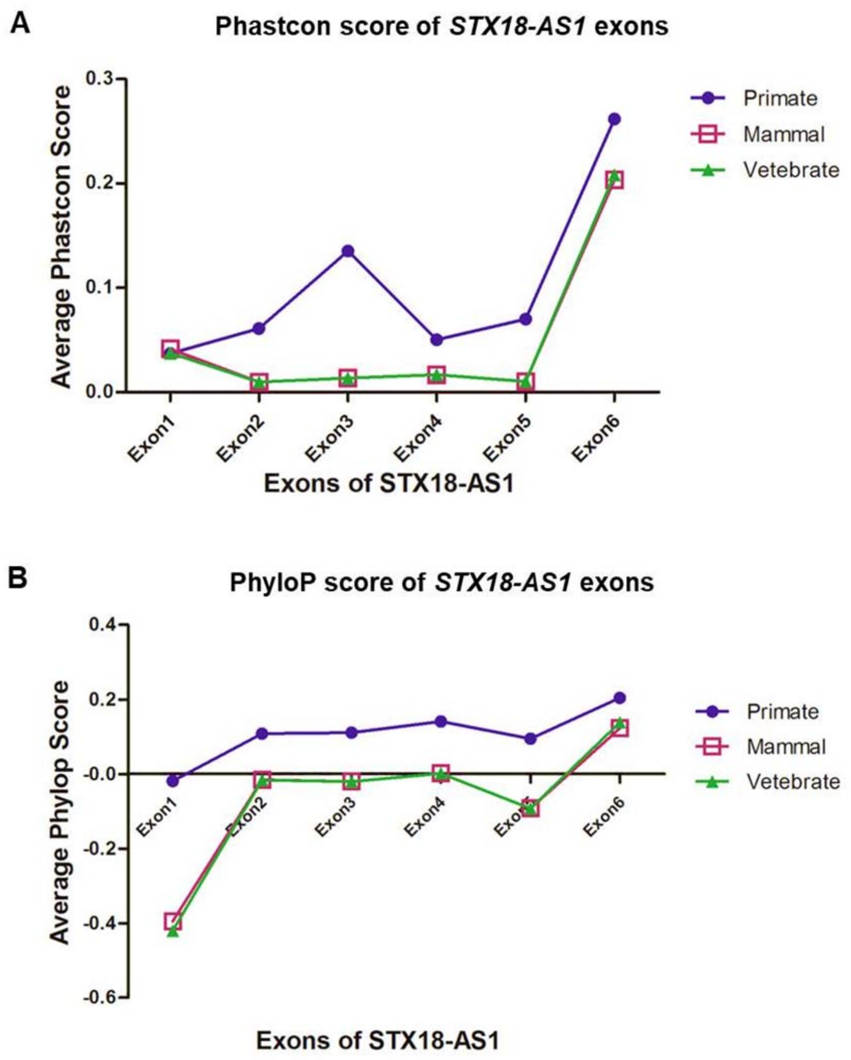


**Supplementary figure 2: The conservation of *STX18-AS1* across species.**

Phastcon (A) and Phylop (B) scores of *STX18-AS1* exons from the UCSC genome browser (https://genome.ucsc.edu/) were collected and compared among Primates, Mammals, and Vertebrates.

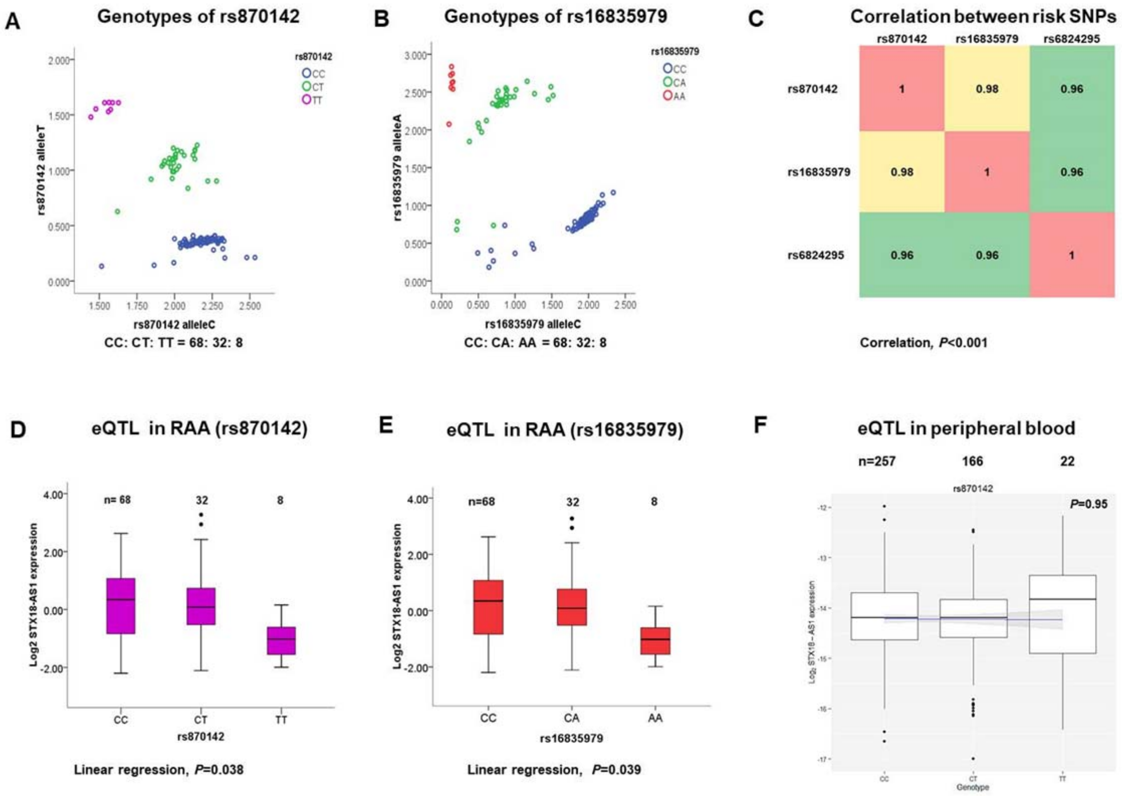


**Supplementary Figure 3: Genotypes of risk SNPs and eQTL analyses in human samples.**

A, genotypes of rs870142. B, genotypes of rs16835979. C, the correlation calculated between all three risk SNPs identified from GWAS. Correlations were statistically significant with $P < 0.001$. D, the linear regression of *STX18-AS1* expression in human right atrial appendages (RAA) referring to genotypes of rs870142. E, the linear regression of *STX18-AS1* expression in RAA referring to genotypes of rs16835979. F, eQTL results achieved with qPCR on 438 RNA samples from human blood. P values for linear regression were shown along with the figures. P= 0.05 (two sided) was adopted as the statistical significance threshold.

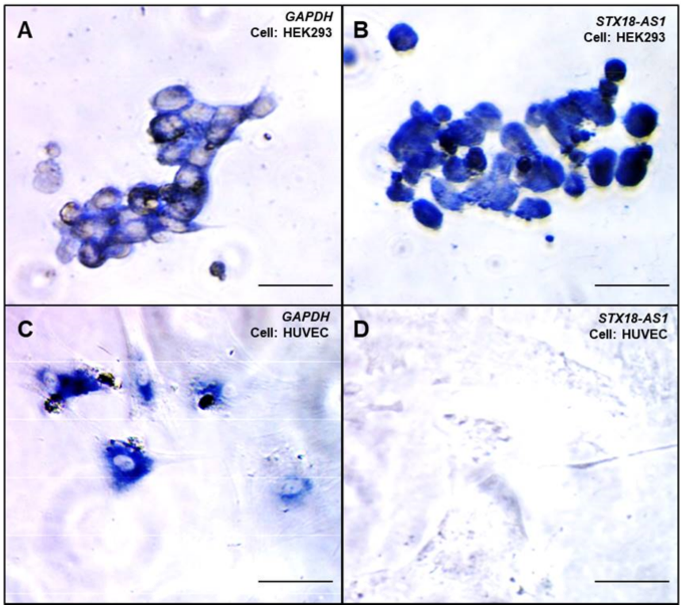


**Supplementary Figure 4: *STX18-AS1* RNA probe evaluation for *in situ* hybridisation.**

*STX18-AS1* is known to be expressed in HEK293 cells (GSM258517 [3]) but at almost undetectable levels in HUVEC cells (https://genome.ucsc.edu/) which serve as a negative control. *GAPDH* was detected in the cytoplasm of both HEK293 **(A)** and HUVEC **(C)** cells. *STX18-AS1* was detected in the nucleus of HEK293 cells **(B)** but not in HUVEC **(D).** Scale bars are 50µm.

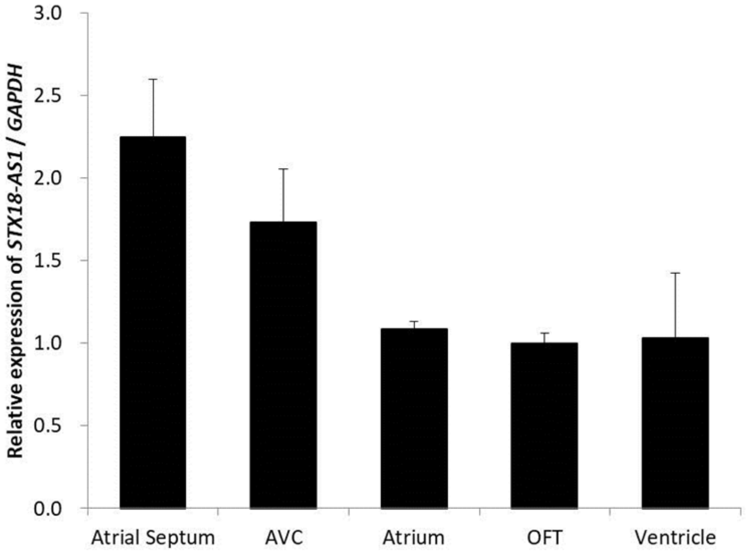


**Supplementary Figure 5: The expression of *STX18-AS1* in different parts of the human embryonic heart.**

The RNA was extracted from the Atrial Septum, Atrioventricular canal (AVC), Atrium, Out-Flow Tract (OFT), and Ventricle of a human embryonic heart of CS15. Only one biological sample was available. Data are shown in Mean±SD (three technical replicates). *GAPDH* was used as the reference gene.

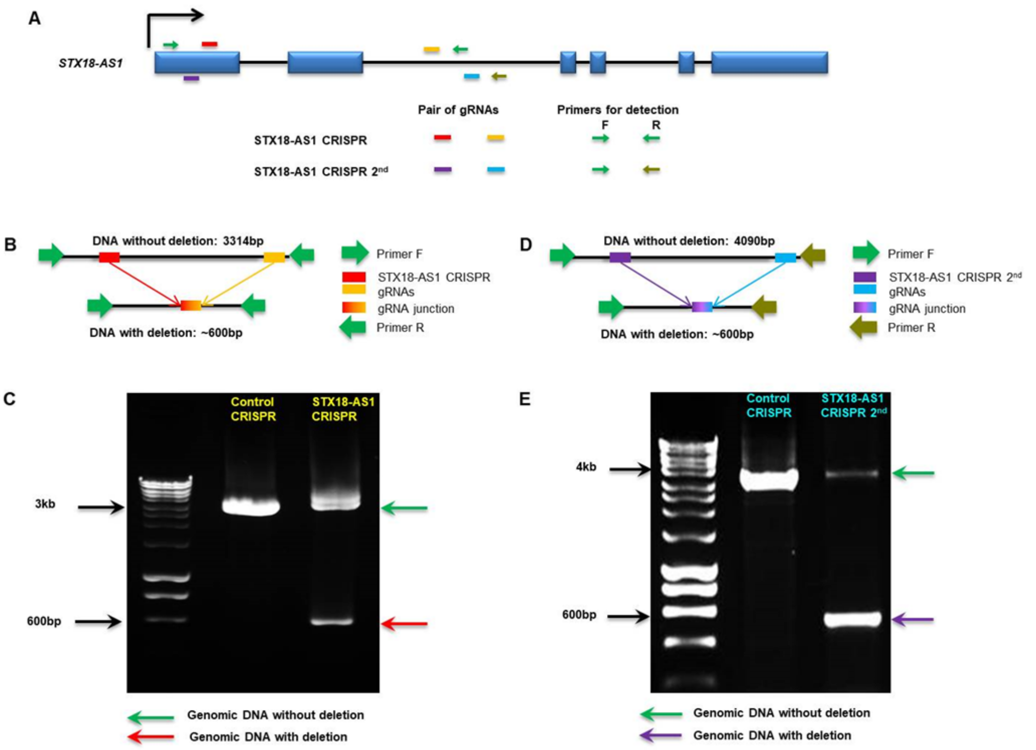


**Supplementary figure 6: CRISPR deletions of *STX18-AS1* in HepG2 cells**.

**A**, designs of CRISPR gRNAs and primers for detection. **B**, the scheme of PCR products from the first design of CRISPR “STX18-AS1 CRISPR”. **C**, PCR results of genomic DNA with STX18-AS1 CRISPR deletions. **D**, the scheme of PCR products from the second design of CRISPR “STX18-AS1 CRISPR 2$^{nd}$”. **E**, PCR results of genomic DNA with knockdown effects of STX18-AS1 CRISPR 2$^{nd}$.

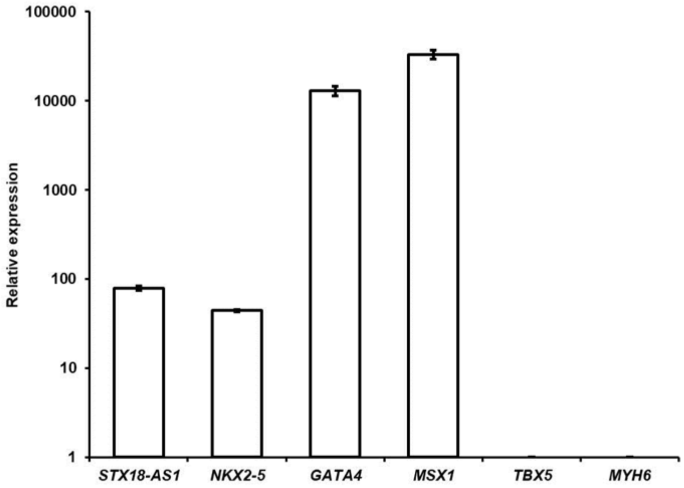


**Supplementary figure 7: The expression of *STX18-AS1* and cardiac genes in HepG2 cells.**

Expression data were analysed in reference to *IPO8* and normalized to *TBX5* (as 1), the CT value of which was 38. *MYH6* was not detected with qPCR. Data are shown in Mean±SD.

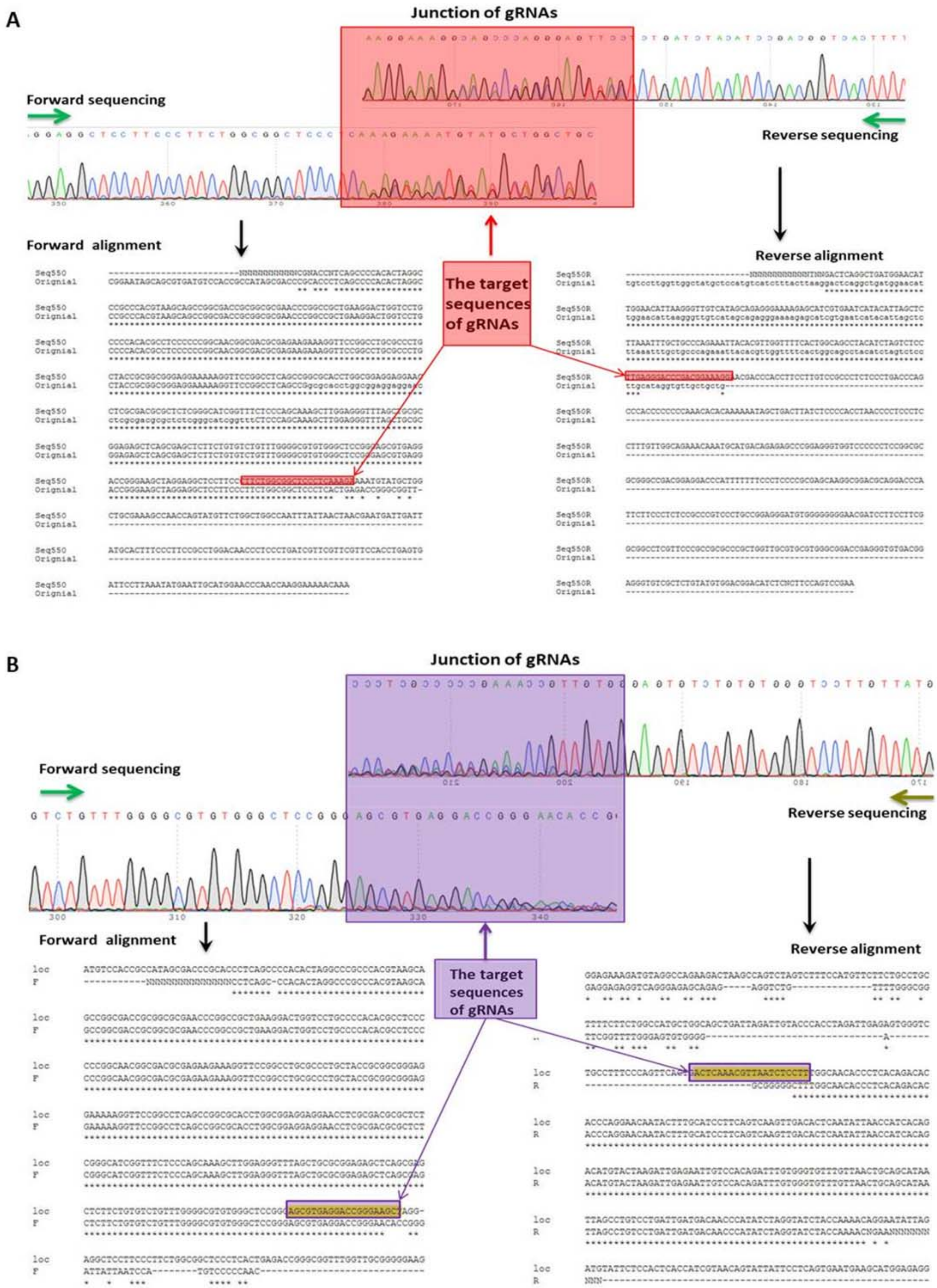


**Supplementary figure 8: Sequence alignment of PCR products with *STX18-AS1* knockdown**

**A**, sequence alignment of PCR products from STX18-AS1 CRISPR with both forward and reverse strands. Red region marking the junction of deletions, also the targeting sequence of gRNAs. **B**, sequence alignment of PCR products from STX18-AS1 CRISPR 2nd from two directions. Purple region marking the junction of deletions and gRNA targeting sequences.

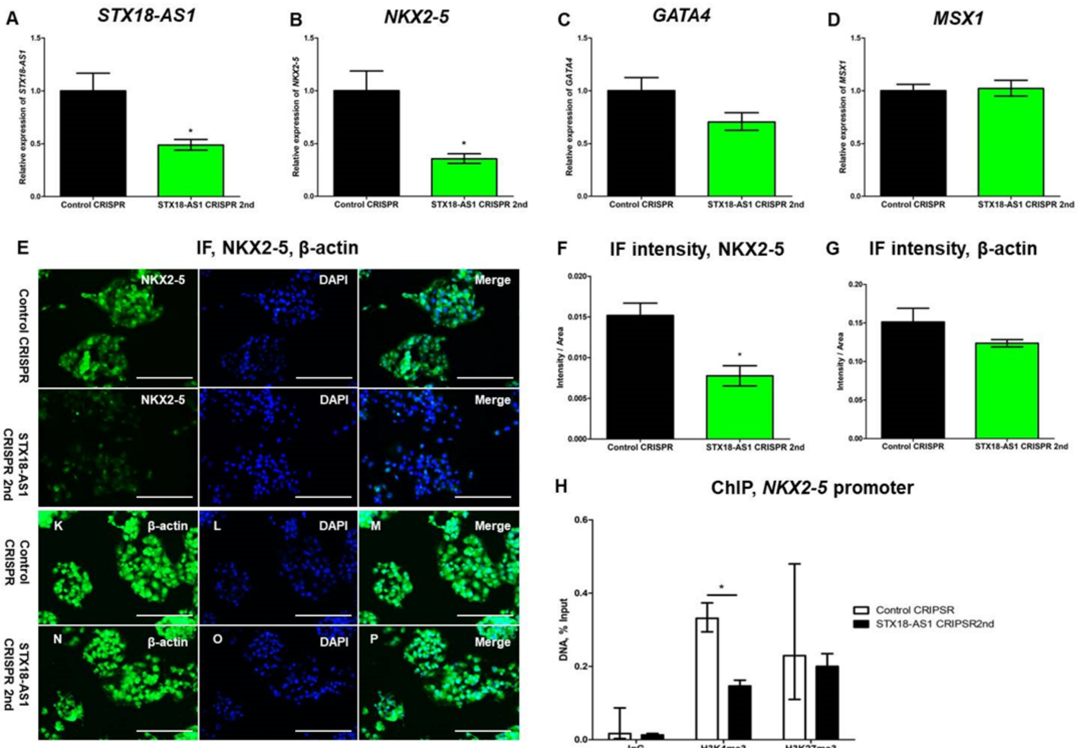


**Supplementary figure 9: Knockdown with STX18-AS1 CRISPR 2nd regulates the expression of *NKX2-5* in HepG2.**

With STX18-AS1 CRISPR 2nd, the transcription of *STX18-AS1* **(A)** and *NKX2-5* were reduced **(B),** without changes in *GATA4* **(C)** and *MSX1* **(D).** The decrease in NKX2-5 protein was detected with immunofluorescence **(E)** comparing with β-actin. The difference between Control CRISPR and STX18-AS1 CRISPR 2nd was statistically significant in intensity analyses **(F-G).** Upon STX18-AS1 CRISPR 2nd knockdown, the histone modification of H3K4me3 around *NKX2-5* was inhibited without changed in H3K27me3 **(H).** Data are shown as Mean±S.E. *, $P<0.05$, comparing with Control CRISPR. Scale bars are 200μm.

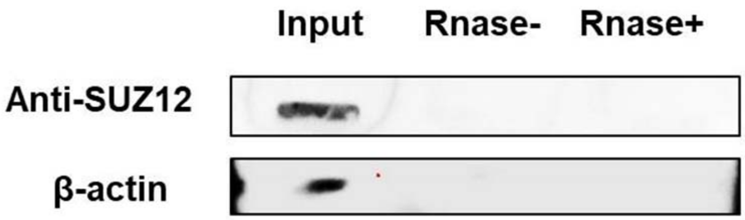


**Supplementary figure 10: Undetected SUZ12 in lysate pulled down by *STX18-AS1* probes**

From the same ChIRP for detection of WDR5, another slot blot was conducted using the same protein sample targeting on SUZ12, the core component of PRC2 complex (which catalyzes the methylation of H3K27). Rnase-, pulldown protein sample; Rnase+, RNase treated negative control; β-actin control for protein precipitation was the same as in Figure 4 D.

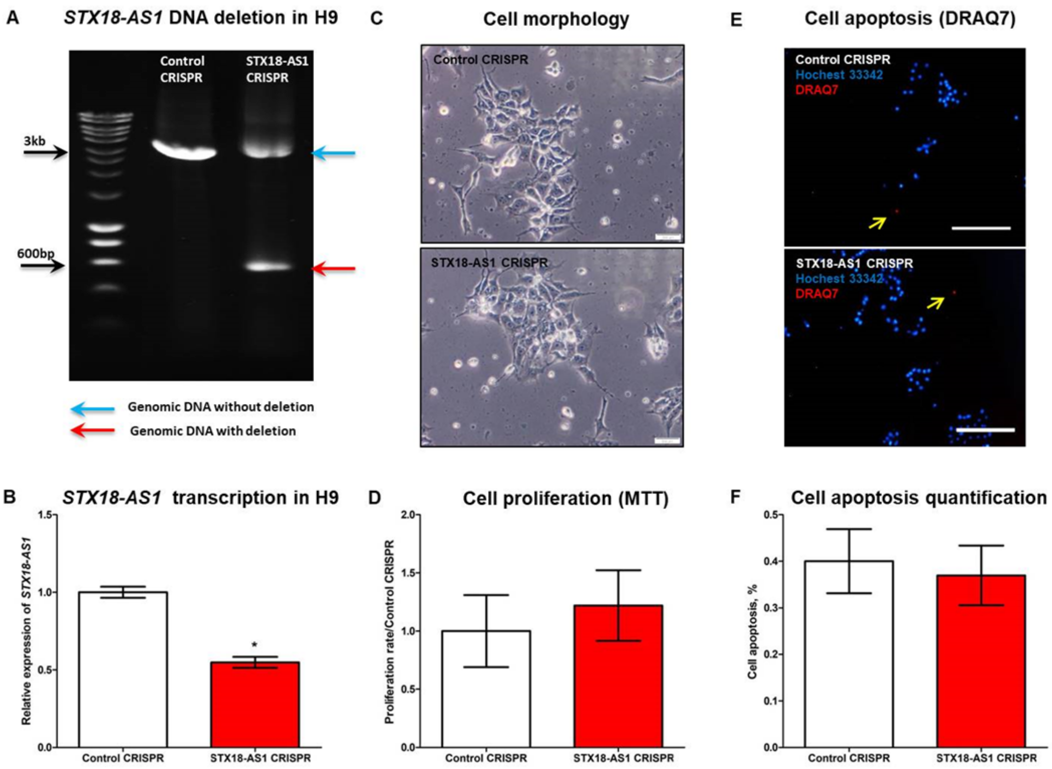


**Supplementary figure 11: Unchanged properties of human embryonic stem cells (H9) with *STX18-AS1* knockdown**

With STX18-AS1 CRISPR, deletions at *STX18-AS1* was achieved in H9 cells at genomic level with a short product of ~600bp **(A),** transcription of *STX18-AS1* reduced accordingly by ~50% **(B).** No visible change was observed in cell morphology **(C),** proliferation rate **(D),** and cell apoptosis **(E-F).** Cells in **E** were double stained with DRAQ7 (apoptosis cells, red colour with yellow arrows) and Hochest33342 (the nucleus of live cells, blue). Data are shown in Mean±S.E. *, *P*<0.05, comparing with Control CRISPR. Scale bars are 100μm for **C** and 200μm for **E**.

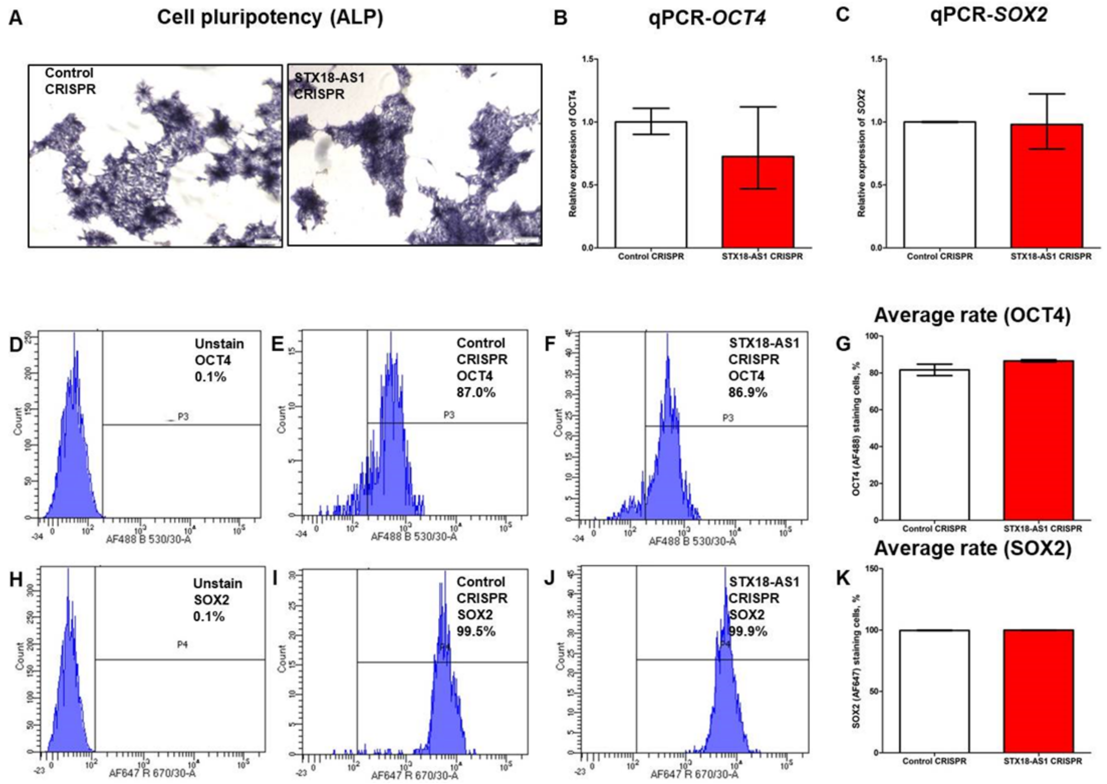


**Supplementary figure 12: Unchanged cell pluripotency of H9 cells with *STX18-AS1* knockdown.**

**A**, Pluripotency detected by ALP assay. **B-C,** unchanged transcription of *OCT4* and *SOX2* (gene markers for stem cell pluripotency [4]) in Control CRISPR and STX18-AS1 CRISPR. **D-F**, FACS analyses with OCT4-AF488 antibody, with "**G**" as the quantitative analysis of cell staining rate. **H-J**, FACS analyses with SOX2-AF647 antibody, with "**K**" as the quantitative analysis of cell staining rate. Scale bars are 500μm. Data are shown in Mean ± S.E. Significant level was sat as $P$<0.05.

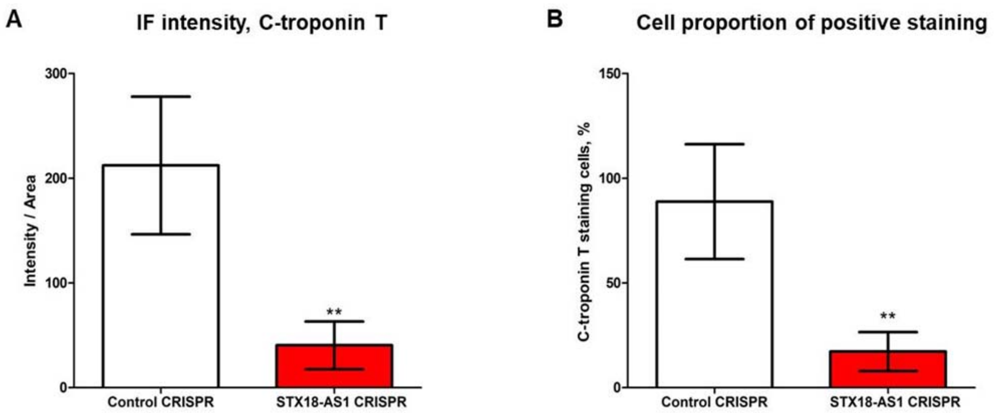


**Supplementary figure 13: Quantification of the changes of C-troponin T staining after *in vitro* cardiomyocyte differentiation.**

H9 cells with Control CRISPR and STX18-AS1 CRISPR were conducted with the same procedures and stained with C-troponin T antibody. Immunofluorescent pictures were taken at 20× magnification. Intensity and numbers of positive staining cells were quantified with ImageJ reference to DAPI. 3 biological replicates for each group. Data are shown as Mean±S.E. *, $P$<0.05, comparing with Control CRISPR.
