## Supplementary file for "*STX18-AS1* is a Long Noncoding RNA predisposing to Atrial Septal Defect via downregulation of *NKX2-5* in differentiating cardiomyocytes"

### Supplementary methods

#### 1. CRISPR gRNA designs

The gRNAs were designed with an online tool developed by Zhang's Lab [1] (http://crispr.mit.edu/) and ATUM grna design tool (https://www.atum.bio/eCommerce/cas9/input). Designs with high scores in both tools were selected for CRISPR construction.

Table S1 Designs of CRISPR gRNA pairs

| **The 1st pair of gRNA, STX18-AS1 CRISPR** | |
|---|---|
| **Upstream gRNA** | AACCGCCCGGTCTCAGTGAGGG |
| **Downstream gRNA** | CAGCAGCAACACCTATGCAAGG |
| **The 1st pair of gRNA, STX18-AS1 CRISPR 2nd** | |
| **Upstream gRNA** | AGCGTGAGGACCGGGAAGCTAGG |
| **Downstream gRNA** | ACTCAAACGTTAATCTCCTTTGG |

Table S2 PCR primers for CRISPR genomic DNA qualification

| **Target DNA deletion** | **Forward** | **Reverse** |
|---|---|---|
| **STX18-AS1 CRISPR** | CGGAATAGCAGCGTGATGTC | tgtccttggttggctatgct |
| **STX18-AS1 CRISPR 2nd** | CGGAATAGCAGCGTGATGTC | actgttacgatggtgagtgga |

#### 2. Assays for qPCR

The genotyping and expression detection in this study was produced by qPCR with TaqMan probe assays. The reference IDs of assays applied were listed in Table S3.

Table S3 TaqMan assays for qPCR:

| **Target** | **Purpose** | **Assay ID** |
|---|---|---|
| **rs870142** | Genotyping | C___8840003_10 |
| **rs16835979** | Genotyping | C__34027652_20 |
| ***STX18-AS1*** | Gene expression | Hs00416742_m1 |

| | | |
|---|---|---|
| ***NKX2-5*** | Gene expression | Hs00231763_m1 |
| ***GATA4*** | Gene expression | Hs00171403_m1 |
| ***MSX1*** | Gene expression | MSXPROB design from [2] |
| ***TBX5*** | Gene expression | Hs00361155_m1 |
| ***MYH6*** | Gene expression | Hs01101425_m1 |
| ***OCT4*** | Gene expression | Hs04260367_gH |
| ***SOX2*** | Gene expression | Hs01053049_s1 |
| ***IPO8*** | Gene expression | Hs00183533_m1 |
| ***GAPDH*** | Gene expression | Hs99999905_m1 |

3. DIG Labeling of RNA probes

The RNA probe for *in situ* hybridisation was designed spanning the first three exons of *STX18-AS1* (514bp). The sequence of the probe was amplified with PCR with a forward primer (GCGAGCTCTTCTGTGTCTGT) and reverse primer (TGCTGGAAGACACAGGCTTT) tagged by T3 sequence (AATTAACCCTCACTAAAGGG). T3 sequence is the recognisable start site in *STX18-AS1* cDNA for *in vitro* transcription with T3 polymerase (Promega) and DIG RNA probe labelling kit (Roche). The *in situ* hybridisation probe of GAPDH (527bp) was designed for intracellular location comparison (forward primer: ACCACCATGGAGAAGGCTGG; reverse primer: CTCAGTGTAGCCCAGGATGC). For the synthesis of sense probe control, the T3 sequence was attached to the forward primer. After transcription, the DIG tagged RNA probes were purified and stocked at -20°C for future use *in situ* hybridisation.

4. Microtomy and paraffin section

Hearts from ISH were dehydrated gradually with 50% Ethanol, 75% Ethanol, 100% Ethanol (30min for each) with the following dehydration step with 100% Ethanol overnight. The histological clearing of hearts was produced with successive incubation (30min) in 50% HistoClear (with Ethanol, RT), 100% HistoClear (RT), and 100% HistoClear (65℃). The hearts were then transferred into paraffin (65℃) overnight for stabilization before embedding into blocks. The chilled tissue blocks were processed to sectioning with Microtome (Leica RM2145). The sections were trimmed with the thickness of 6-8 μm and flattened on the surface of pre-warmed water (42℃). The picked up sections by slides were dried at 42℃ overnight. The slides with sections were stained with Eosin after dewaxing with Xylene and rehydration in water. In the final, the slides were mounted with DPX Mounting Media (Merck) for observation.

5. Biotin tagged *STX18-AS1* probes for ChIRP

Following the protocol of Chu et al's [3], the probes for ChIRP were designed at http://singlemoleculefish.com/ with about one probe per 100bp (20bp in length, 45% GC, spacing length 60-80, omitting repeat sequences). For the full transcript of *STX18-AS1* (ENSG00000247708), in total 15 probes were designed in the following. All probes were biotin labelled. The designs of positive control probe U1 (ctcccctgccaggtaagtat) and negative control probe (ccagtgaatccgtaatcatg) are adapted from Chu et al's study [4].

Table S4 ChIRP probes for *STX18-AS1*

| Probe ID | Sequence |
|---|---|
| STX18-AS1_1 | tttgctgggagaaaccgatg |
| STX18-AS1_2 | gaagggaaggagcctcctag |
| STX18-AS1_3 | tctatcctgctcaaggtgag |
| STX18-AS1_4 | cccacatgatgaaagctgat |
| STX18-AS1_5 | ccttgttagtgtatgaaggc |
| STX18-AS1_6 | tggcaatgacatttaccagc |
| STX18-AS1_7 | acagctaaggcttttcacac |
| STX18-AS1_8 | tctgtagtgcttggagagat |
| STX18-AS1_9 | agtgtggccattcattaaga |

| | |
|---|---|
| STX18-AS1_10 | ggtctcaccaaagaaatggc |
| STX18-AS1_11 | agtggcggaattgagtctta |
| STX18-AS1_12 | gtctcaagtgggttttctgg |
| STX18-AS1_13 | ttccttgacatttcaatcct |
| STX18-AS1_14 | tctctcttacaaagcatact |
| STX18-AS1_15 | cactggttacattttccgac |

6. Trichloroacetic Acid (TCA) protein precipitation

For each 1ml of eluent from ChIRP, 250μl TCA was added for protein precipitation at 4℃ overnight. At the next day, samples were centrifuged at 16000g for 30min (4℃) to pellet proteins. The pellets were washed with cold Acetone for two times before air drying (1min). The protein pellets were then dissolved in 1×Laemmli sample buffer and boiled at 95℃ (30min) for immunoblotting or slot blotting.

7. Fluorescence-activated cell sorting (FACS) for pluripotency detection

For FACS analyses with intracellular markers, cells ($10^{\wedge 6}$/sample) were firstly crosslinked with 3% formaldehyde for 20min before antibodies incubation. Cells were blocked with 5% FBS in PBS (flow buffer) for 30min after three PBS washes. In the following, cells were resuspended in 100ul flow buffer and incubated with antibodies, AFS647-SOX2 (1μl, Biolegend) and AFS488-OCT4 (5μl, Biolegend) for 1 hour at 4□ in the dark. After another three washes with flow buffer, cells were collected and resuspended in 500ul flow buffer for FACS reading. Unstained cells were used as background controls for analyses.
